# Explicit and implicit depth-cue integration: evidence of systematic biases with real objects

**DOI:** 10.1101/2021.03.19.436171

**Authors:** Carlo Campagnoli, Bethany Hung, Fulvio Domini

## Abstract

In a previous series of experiments using virtual stimuli, we found evidence that 3D shape estimation agrees to a superadditivity rule of depth-cue combination. According to this rule, adding depth cues leads to greater perceived depth magnitudes and, in principle, to depth overestimation. The mechanism underlying the superadditivity effect can be fully accounted for by a normative theory of cue integration, through the adaptation of a model of cue integration termed the Intrinsic Constraint (IC) model. As for its nature, it remains unclear whether superadditivity is a byproduct of the artificial nature of virtual environments, causing explicit reasoning to infiltrate behavior and inflate the depth judgments when a scene is richer in depth cues, or the genuine output of the process of depth-cue integration. In the present study, we addressed this question by testing whether the IC model’s prediction of superadditivity generalizes beyond VR environments to real world situations. We asked participants to judge the perceived 3D shape of cardboard prisms through a matching task. To assay the potential influence of explicit control over those perceptual estimates, we also asked participants to reach and hold the same objects with their fingertips and we analyzed the in-flight grip size during the reaching. Using physical objects ensured that all visual information was fully consistent with the stimuli’s 3D structure without computer-generated artifacts. We designed a novel technique to carefully control binocular and monocular 3D cues independently from one another, allowing to add or remove depth information from the scene seamlessly. Even with real objects, participants exhibited a clear superadditivity effect in both explicit and implicit tasks. Furthermore, the magnitude of this effect was accurately predicted by the IC model. These results confirm that superadditivity is an inherent feature of depth estimation.

## Introduction

In previous studies we showed that, when sources of depth information (cues) are added to the virtual rendering of a 3D shape, the perceived magnitude of a given 3D property of that shape (e.g., its depth, slant or curvature) increases. For instance, when viewing a cylindrical surface from a close distance (50 cm), its overall depth is consistently underestimated when the shape is only specified by binocular disparity, compared to when specified by a mixture of disparity signals and texture gradient (Campagnoli & Domini, 2019).

This pervasive phenomenon, which we refer to here as superadditivity effect (the estimated depth from a combination of depth cues is greater than the depth estimates from the individual cues) has been previously observed in the context of a series of perceptual studies investigating a model of depth cue combination termed Intrinsic Constraint (IC) model (Di Luca et al., 2010; Domini & Caudek, 2003, 2009, 2010, 2011, 2013; Domini et al., 2006; Domini et al., 2011; Kemp et al., 2018). The IC model can be adapted so to successfully predict these kinds of shifts in 3D shape estimation, by pooling the retinal depth cues through a combination rule that prioritizes maximizing sensitivity to depth information over accuracy (Domini & Vishwanath, 2020).

While the model provides a description of the mechanism behind the behavioral effect, it is unclear whether the presence of superadditivity in the first place is the symptom of an explicit overcompensation introduced by the observers, or the signature of a characteristic feature of depth estimation. On the one hand, when witnessing more depth signals in the visual field, superadditivity may be the result of participants trying to qualitatively express the greater richness of their visual experience by consciously inflating their judgments. If, on the other hand, superadditivity relates directly to the readout of the internal depth estimates, we should expect it to equally affect non-explicit tasks such as reaching towards a 3D object with the hand to pick it up. Converging evidence in support of the latter possibility has been documented in previous VR investigations (Foster et al., 2011; Campagnoli & Domini, 2016, 2019). However, the artificial setups used in those studies may have introduced yet another potential source of explicit bias in the responses, because participants immersed in a virtual scene tend to supervise their behavior and exert top-down control over their own actions (Chessa et al., 2019).

To investigate the nature of the phenomenon, we sought to test the predictions of the IC model for the superadditivity effect in the context of a real-world scene, by asking participants to judge as well as to interact with physical objects. To have the same visual conditions of the previous VR experiments, we designed a novel technique to manipulate the amount of disparity, motion and focus cues specifying the shape of a cardboard-made prism, and we examined how participants judged the stimuli’s shape under the various visual conditions. In a within-subjects design, we compared explicit reports of perceived depth with an implicit measure of depth provided by the in-flight grip aperture when participants reached with the hand towards the same object to hold it. As a generalization test of the predictions, the stimuli were purposely different in shape from those used before. The results were qualitatively identical to what previously obtained in VR, supporting the hypothesis that the visuomotor system integrates depth cues following a combination rule that is not triggered by computer-generated environments, but describes a more general principle of how 3D shapes are represented internally by the brain.

### The mechanism of superadditivity predicted by the IC theory of cue integration

A traditional assumption in modeling cue integration is that separate visual modules analyze independent sources of information to derive magnitudes of distal 3D properties (e.g., the overall depth of an object, the slant of a plane, the curvedness of a smooth surface) in an accurate fashion. Although the assumption of veridicality is lawful both theoretically and computationally, it is often challenged by empirical findings showing that 3D shape estimation is affected by systematic distortions under a variety of visual conditions (Glennerster et al., 1996; Hecht et al., 1999; Tittle et al., 1995; Tassinari et al., 2008; Todd & Norman, 2003). One way to reconcile this evidence is through the post-hoc introduction of additional free parameters to the model – for example prior terms accounting for specific biases in the data – with the downside that they may potentially lead to overfitting. Another way is by seeking a combination rule that aims to maximize something other than accuracy, without increasing its complexity.

The IC theory of cue integration follows the latter approach and posits that the goal of the visual system is to maximize the sensitivity to the available 3D information, while minimizing the undesired influence of other scene parameters. According to this framework, independent visual modules provide 3D estimates that are only linearly related to the magnitude of physical 3D properties, not veridical to begin with. Veridicality is still allowed under ideal conditions, but as a special case of the theory. In general, a given 3D estimate 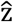 is assumed to be proportional to the corresponding distal 3D property z, through a proportionality factor that expresses the strength of the visual information at any given time.

To illustrate the concept of information strength in a visual scene, let’s consider how depth is estimated from two fundamental cues: binocular disparity and motion. At a distance within the peripersonal space of an agent, depth judgements from relative disparity are usually quite accurate. Formally, we can re-write this statement as 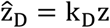, with k_D_= 1. When objects are farther away from the observer, certain distance information that is necessary to scale binocular disparities becomes weaker (k_D_< 1). Eventually, at very large distances k_D_approaches zero. In a similar fashion, depth from motion information depends on the relative motion between the observer and the distal object. For ideal conditions, 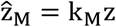 with k_M_= 1. However, for a static observer the proportionality factor k_M_is nil.

In other words, separate 3D modules analyze distinct depth cues (e.g., disparity and motion) which are affected by independent confounding variables associated with viewing conditions (e.g., fixation distance and observer’s motion, respectively). For the i-th depth cue, the parameter k_i_embodies the effect of those confounding variables on the mapping between the physical depth z and its estimate 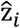. In the framework of the IC theory, we will refer to k_i_as cue strength.

Goal of the IC model is *to maximize sensitivity to depth while keeping the influence of confounding variables to the minimum possible*. In the Appendix (Proof 1) we show that this goal is achieved computationally by a model that combines single-cue estimates through a vector sum. For the general case of a vector space with n image signals, the combined estimate 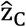 is the resultant vector given by the equation:

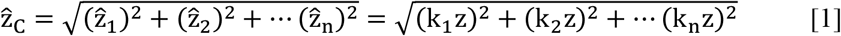

Note that this model is additive, that is, increasing the number of cues yields larger depth estimates. This should not be misunderstood as the combination rule producing systematic overestimations of depth as more cues are added to a stimulus. On the contrary, the underlying logic is that removing cues brings the model outside of its optimal operating conditions, which results in the underestimation of single-cue stimuli (namely, stimuli isolating only one source of information).

The goal of this study was twofold. First, it aimed to investigate whether the phenomenon of superadditivity observed in VR generalizes to the physical world and is therefore inherent to the process of depth-cue combination. To this end, we used real objects and measured their estimated depth through both an explicit (perceptual) and an implicit (motor) task. Second, the study also served as a quantitative test of the IC model predictions. This second aim requires a careful assessment of all individual depth signals that are at play in a given visual scene. In principle, to isolate a specific depth cue and be able to measure its estimate 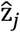, one must reduce the strength k_i_of all the other cues to zero, such that the combined estimate 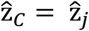 as per equation [1]. This logic applies to both VR and real-world environments. For instance, a static object does not provide any depth from motion information – which in IC terms means that the strength of the motion cue k_M_is equal to zero. Certain types of depth signals, however, cannot be eliminated, and that is because they do not depend on the visual input itself but on the limits of the human optical apparatus. We will use the term “focus cues” to refer to all these signals at once, of which the most relevant one is the blurring gradient. In a VR setting, focus cues specify the flat surface of the monitor screen (z = 0), in contrast with the three-dimensional quality of the simulated environment. With real objects, focus cues add 3D shape information on top of, and consistent with, what already specified by stronger depth cues such as disparity or motion.

Since in the present study we investigated the contribution of stereo and motion cues to the estimation of depth, equation [1] becomes:

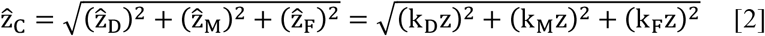

where the depth estimates 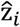 are all related to the same physical object’s depth z, but each is scaled by the strength term appropriate for the signal in question (k_D_, k_M_and k_F_ for disparity, motion and focus cues, respectively). Importantly, focus cues are dramatically different in computer-generated versus real-world scenes. VR environments require the presence of a flat screen, therefore the depth associated with the focus cues is zero and the last term of equation [2] is dropped. Under these visual conditions, it is possible to study a true disparity-only (that is, motion has strength k_M_= 0) condition and test whether adding motion to the display (k_M_≠ 0) yields larger magnitudes of perceived depth. This was done in our past study (Campagnoli and Domini, 2019). Consider now a real object viewed at a distance yielding sizable focus cues, like a reachable distance. Unlike VR, this viewing condition does not enable to isolate disparity and motion cues, because the focus cues are also attuned to the 3D structure of the same object, given that the scene is no longer simulated on a flat screen. As a result, the “disparity-only” stimulus will in fact be a combination of disparity and focus signals (yielding a depth estimate 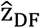), and the same will be true for the “motion-only” stimulus (yielding a depth estimate 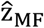). This will impact how the combined estimate of depth 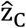 is derived from the three cues. It can be demonstrated (see Appendix, Proof 2) that combined cue estimates from “single-cue” estimates is given by:

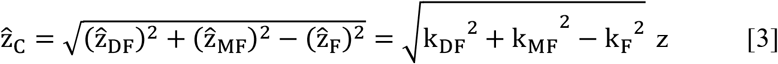

This operational definition allows to make specific quantitative predictions about superadditivity, since equation [3] is effectively expressed as a function of “single-cue” estimates. In this investigation, we tested this prediction of the IC model with stimuli that allowed to equalize the strength of the disparity and the motion cues (k_S_≈ k_M_). This way, we expected that the psychometric functions relating disparity-and motion-derived depth estimates to the physical depth of the stimuli would have approximately the same slope (Fig. 1, red and blue lines). Secondly, we also expected a significant effect of the focus cues (k_F_≠ 0, green line in fig. 1). Finally, equation [3] allowed to predict the slope of the combined-cue psychometric function (Fig. 1, black line) without introducing additional free parameters, but only from the slopes (strength terms) of the single-cue functions.

**Figure 1.**
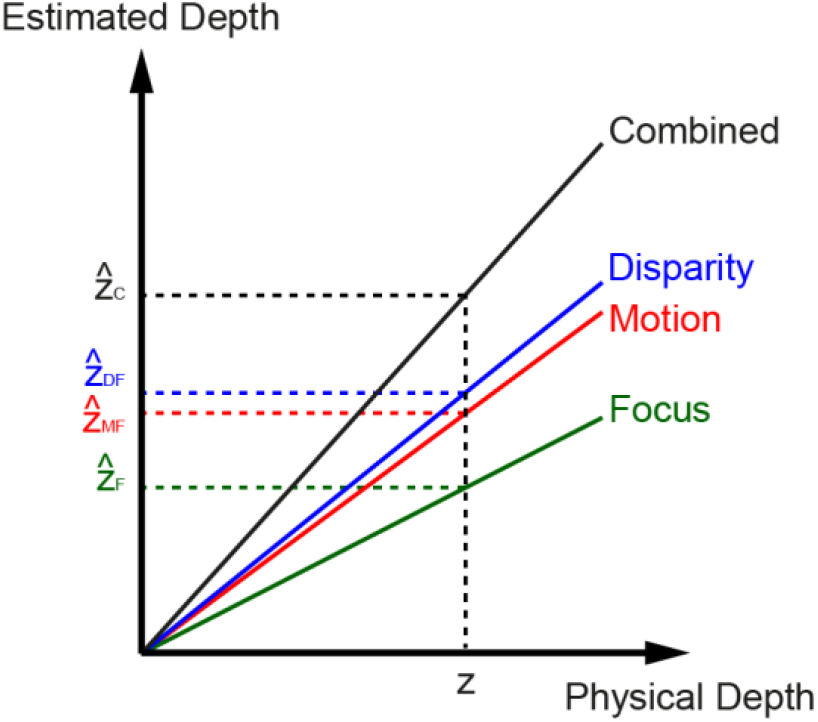
Superadditivity prediction: estimates of depth (values on the ordinate) were expected to be greatest for the visual condition with the maximum amount of 3D sources (Combined, in black), intermediate under Disparity and Motion (blue and red respectively), and smallest for the condition with the scarcest amount of depth information (Focus, in green). The increase in slope of the Combined estimate was predicted on the basis of the slope of the other conditions according to equation [3].

### Manipulating depth information with real objects

In a natural environment, binocular vision and motion parallax are among the most important depth cues available in the visual array. While it is relatively easy to balance depth cues in VR, it is much harder to do so in a natural environment. The biggest challenge is represented by stereovision, since objects seen within reaching distance inevitably produce a strong stereo signal which prevails over the other signals to depth. On the other hand, the pattern of retinal velocities resulting from the relative motion between object and observer can be easily modulated by rotating the object at different speeds.

To overcome this problem, we designed a novel technique to allow binocular disparities to have similar strength to that of retinal velocities for the judgment of depth, without using optical devices or introducing other artificial surrogates.

We constructed prism-shaped objects out of cardstock, to be used as stimuli (Fig. 2A). When oriented horizontally, the visible portion of the stimuli consisted of 60 mm wide, 40 mm tall and 20 or 40 mm deep wedge. The surface of the objects was matte and painted with white stripes that ran parallel to the long side. Subjects saw the visible portion of the prisms in six configurations (Fig. 2B): static horizontal (Focus condition), static tilted (Disparity condition; 10 and 20 deg tilt), horizontal with motion (Motion condition), and tilted with motion (Combined condition; 10 and 20 deg tilt).

**Figure 2.**
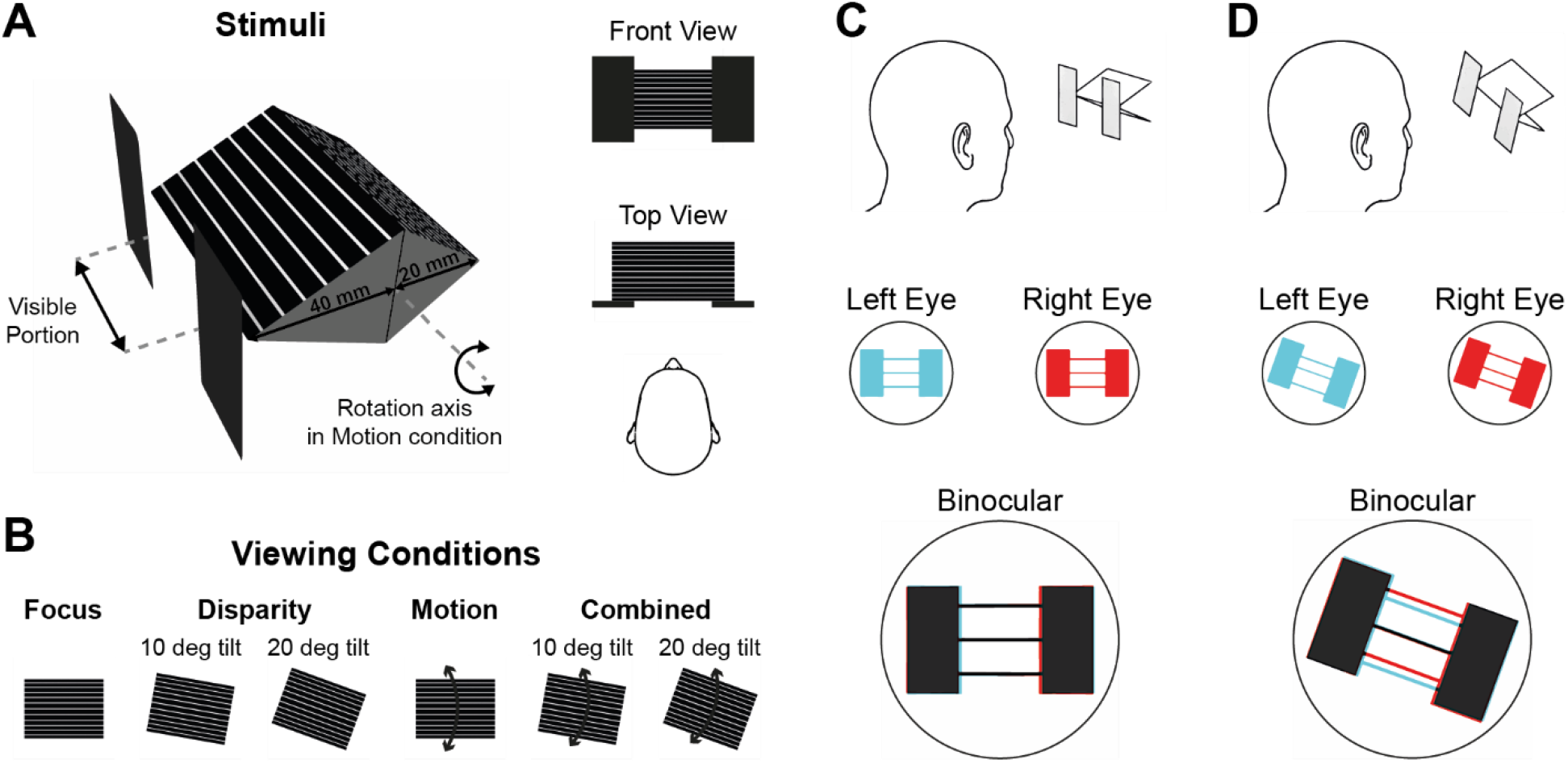
(A) Schematic of the stimuli used in the experiment (see also the pictures in figure 3). Two prisms (20 and 40 mm in depth) were attached back-to-back and displayed one at the time by rotating them 180 degrees around the axis passing through the common face (the same axis for the rocking of the Motion condition). Two rectangular occluders prevented the vision of the edges of the stimulus. (B) Viewing conditions of the experiment. Binocular disparity was gradually added by tilting the stimulus relative to the frontoparallel plane (Disparity). Motion was added by rocking the stimulus around its main axis at constant angular velocity. (C) When the stimulus is horizontal, the left and right retinal projections (in cyan and red, respectively) align such that after fusion (“Binocular” inset) the binocular disparities from the prism’s surface are unspecified, because the correspondence problem cannot be solved. (D) Tilting the stimulus relative to the frontoparallel plane disambiguates the binocular disparities: note the greater binocular disparity (misalignment of the cyan and blue projections) at the top and bottom edges of the prism relative to the tip (assumed to be at fixation). The misalignment of the retinal projections increases up to a maximum tilt angle of 90 degrees (vertical stimulus – see Van Ee & Schor, 2000).

During the Focus condition, the prism was static and oriented horizontally along its long side, with the tip facing the observer. When in this position, the horizontal disparities were unspecified, since the uniformity of the horizontal lines on its surface made the correspondence problem unsolvable (figure 2C). The overall angular size of the visible stimulus (about 8 degrees) was such that vertical disparities were negligible. Moreover, the presence of cardboard occluders eliminated contour information from the stimulus edges. In this visual condition, subjects could rely mainly on focus cues of extraretinal origin, such as ocular vergence and lens accommodation.

In the Disparity condition, the object was static and also tilted around the axis perpendicular to the frontoparallel plane, pivoting around the cyclopean eye. Rotating the surface in a direction orthogonal to the orientation of the stripes provided an efficient way to selectively manipulate the strength of stereo information (for a detailed analysis of the mechanisms involved see Van Ee & Schor, 2000). As the tilt increased, the binocular disparities blended less and less into the white stripes, becoming more resolvable (figure 2D). The limit case would be represented by a tilt of 90 degrees, corresponding to a perfectly vertical prism, yielding maximum binocular disparity. For this study, we selected two small tilt angles, 10 and 20 degrees. This fulfilled two goals: 1) to test if the strength of stereo did increase with a greater tilt angle – as a general test of the technique itself; 2) most importantly, to limit the overall strength of stereo information relative to motion (when present), so that the two signals could contribute to depth perception equally. Overall, the total depth information in the Disparity condition was the sum of the focus cues plus the disparity gradient.

When the stimulus was horizontal and rocking around the horizontal axis (Motion condition), a pattern of relative velocities of the white stripes allowed the observer to judge the prism’s 3D layout from the retinal motion gradient. Note that this rotation did not interact with the gradient of binocular disparities, which remained unspecified. Therefore, the total depth information in this condition resulted from the combination of focus cues plus the motion gradient.

Finally, in the Combined condition, the prism was tilted relative to the frontoparallel plane as in the Disparity condition, and it also rocked along its tilt axis, so that the movement did not change the orientation of the stripes, as in the Motion condition. During this presentation, the binocular disparities generated from tilting the object were combined with the motion gradient produced by the object’s oscillation. This visual condition provided the maximum amount of depth signals specifying the stimulus geometry: focus cues plus disparity and motion gradients.

## Materials and Methods

### Participants

Forty-three students gave written informed consent to participate in the experiment (29 females, age ranging between 18 and 24). All participants self-reported normal or corrected-to-normal vision. The participants received either a reimbursement of $8/hour or coursework credit for their effort. The experiment was approved by the Institutional Review Board at Brown University. Each participant gave informed consent prior to the experiment.

### Apparatus

Subjects sat with their head locked on an optometric chin and forehead rest, installed along one of the short sides of a rectangular table, directly facing the physical stimulus (figure 3A). The stimulus consisted of an isosceles triangular prism made from cardstock, oriented with the tip facing the participant. We built two of these prisms, one with a depth (tip-to-base extent) of 20 mm (small) and the other of 40 mm (large). The prisms were attached base-to-base such that only one was visible per trial (figure 2A). Printed onto the prism surface were pure white pinstripes on a matte black ink base, spaced 6.5 mm apart.

**Figure 3.**
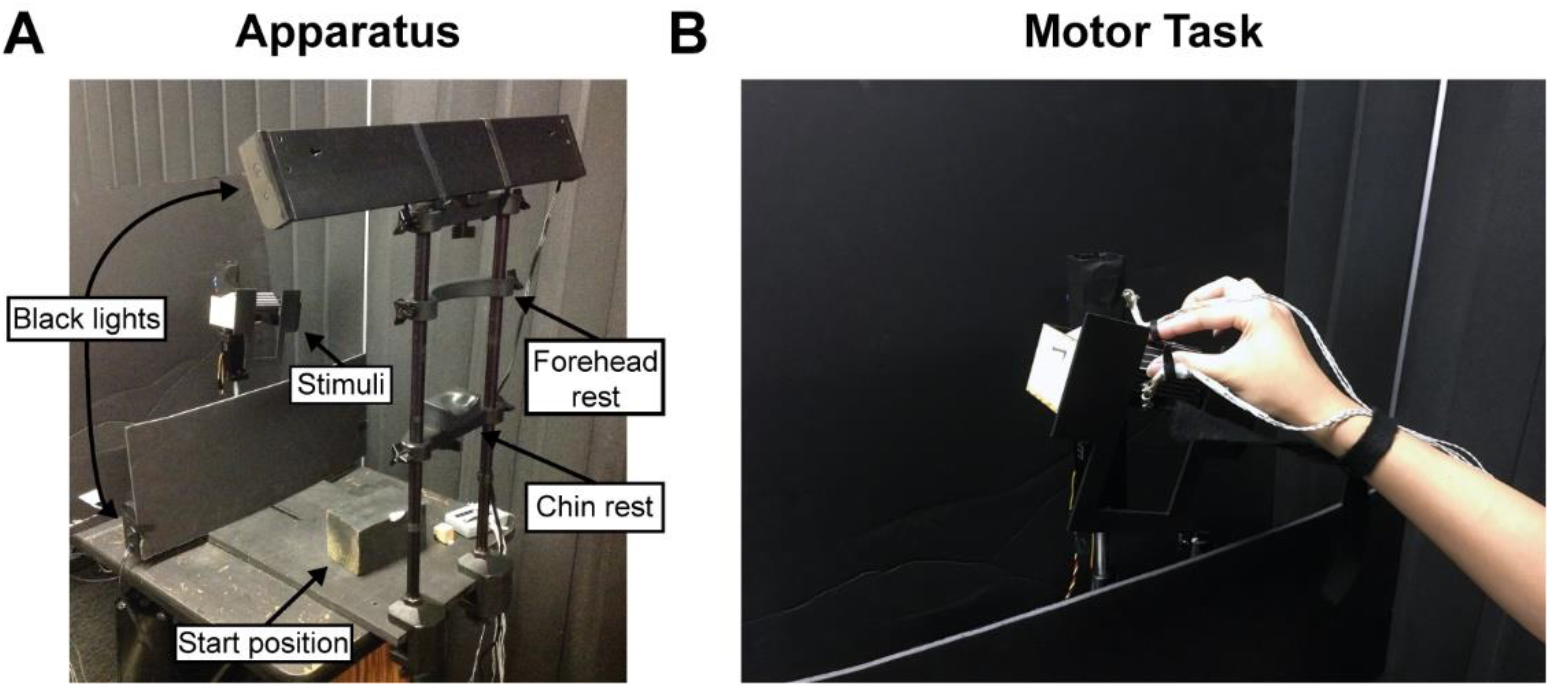
Apparatus (A) and grip gesture in the motor task (B), pictures taken from the actual experimental setup.

During the experiment, the testing room was dark and two black lights guaranteed uniform illumination of the presented stimulus; the net effect was a series of bright white stripes on a dark surface. The two prisms were mounted on a custom, laser-cut wooden base, which was then installed on a vertical metal stand. The position and height of the stand was adjusted such that the tip of the prism was 42 cm above the tabletop, matching the average eye height of the subjects, and either 44 or 42 cm away from the observer, for the small and the large object respectively. Rectangular wooden occluders, also painted matte black, covered the far left and right sides of the stimulus to prevent participants from detecting the edges of the prisms. The visible portion of the stimulus thus was 60 mm wide by 40 mm tall.

Two stepper motors (Phidgets Inc.) were used to switch between stimuli as well as to manipulate their orientation and motion. The first motor rotated the prisms around the horizontal axis centered on their plane of contact (the prisms’ bases), producing an up-and-down rocking motion of the prism’s tip at 60 deg/sec that swept a total angle of 15 degrees (±7.5 degrees, figure 2A and 2B). The second motor rotated the stimulus and occluders around the axis perpendicular to the subject’s frontoparallel plane and centered on the cyclopean eye, setting a total of three angles: 0 degrees (Baseline and Motion conditions), ±10 and ±20 degrees (Disparity and Combined conditions). The values of the angles for the Disparity condition and of the angular velocity for the Motion condition were chosen based on pilot data in order to equalize the strength of stereo and motion information.

Although the integration of disparity and motion cues may lead to two additional sources of binocular information, namely changing disparity over time (CDOT) and interocular velocity difference (IOVD – Harris et al., 2008), these were prevented in the Combined condition. CDOT and IOVD are informative about motion-in-depth, that is when an object is approaching the observer along the line of sight. Our setup ensured that motion-in-depth was negligible. Given the small rotation of the prism about an axis parallel to the white stripes, the motion of each stripe was almost entirely in the front parallel plane, with a negligible component along the line of sight. Consequently, binocular disparities did not change as a result of the rotation, effectively stripping away this potential source of information.

Prior to each trial, the black lights turned off. The first motor rotated to an intermediate position between the prisms, so that subjects could not anticipate which prism was being prepared based on auditory cues, and then positioned the next stimulus. The scene was then illuminated again, via a custom *USB* relay that synced the lights’ power source to the motion of the motors, allowing participants to see the stimulus (and their hand, in the case of the implicit task).

To record the location of the subject’s fingers, we used an *NDI* Optotrak Certus motion capture system, with the position sensor facing the workspace at a distance of 224 cm from the subject. Subjects wore index and thumb markers with protruding flags that held three *NDI* infrared *LEDs* each, oriented such that the infrared light would directly hit the Optotrak cameras. Optotrak recordings, the motion of both Phidgets motors and the *USB* relay were all controlled by custom C++ programs using specific *API* routines provided by each vendor.

### Procedure

Each subject underwent thorough calibration upon putting on the infrared *LEDs*. First, the experimenter calibrated the XYZ axes of the Optotrak position sensor so that the origin was aligned with the observer’s cyclopean eye and the forward direction coincided with the line of sight. Second, the experimenter recorded the position of the subjects’ fingers on a designated home position, marked by a wooden block positioned on the tabletop about 35 cm away from the stimulus (figure 3A). Participants performed an explicit (perceptual) task and an implicit (grasping) task in 4 separate blocks, following an ABBA order.

In the explicit task, the stimulus was illuminated at the beginning of each trial, and subjects performed a Manual Estimation (*ME*), matching the perceived tip-to-upper-edge distance to the proprioception of the distance between thumb and index fingers. The contact surface had length of 28.3 and 44.7 mm for the small and large prism respectively. Subjects were not allowed to look at the hand while performing the task. To confirm their estimate, they pressed a button with the other hand. At the button press, the lights turned off, and the next trial would load as the subjects returned to home position.

In the implicit task, once the lights turned on, subjects reached out and physically contacted the tip and upper rear edge of the stimulus with their thumb and index fingers, respectively (fig. 3B). Subjects could see their hand and feel the prism. If the prism was moving, it stopped once the index markers passed 40 cm in the z direction, to make certain that subjects could firmly contact it. To ensure a comfortable grasping posture, we did not constrain the thumb and index final positions to specific locations along the edges of the prism. On average, subjects approached the object by gradually rotating the wrist, such that the index’s contact position was 8-15 mm farther to the right than the thumb’s contact position. This behavior produced final grip apertures that were systematically larger than the actual stimulus’s side. When subjects returned their fingers to the home position, the lights went off, and the motors prepared the next presentation.

Each task was comprised of 144 trials, divided in two blocks: in each block, a total of six viewing conditions (Baseline, Disparity 10 deg tilt, Disparity 20 deg tilt, Motion, Combined 10 deg tilt, Combined 20 deg tilt) were presented six times for each depth (total of 72 trials). The trials were pseudo-randomized within each block: participants performed six adjacent strings of 12 unique trials (6 viewing conditions times 2 depths) randomized within repetitions. Including both explicit and implicit task, participants performed a total of 288 trials.

### Analysis of kinematic data

Offline analysis of kinematic data was performed using R (R Core Team, 2020). In every trial of every subject, the frame-by-frame 3D positions of index and thumb markers were checked for missing frames (replaced through spline interpolation), smoothed with a fourth-order Savitzky-Golay filter, and used to compute the instantaneous Euclidean distance between them, or Grip Aperture (*GA*). The thumb path was then segmented into regular intervals (or bins) of 2 mm each, and the mean GA was calculated in each bin of each trial of each subject. The GA’s grand mean and variance were then calculated by pooling across the trials of each viewing condition and object side (bin-by-bin, subject-by-subject).

Further dependent variables extracted from the trajectory data (i.e., Maximum Grip Aperture) were analyzed using standard tests, such as repeated measures ANOVA and t-test.

## Results

In this study, we sought to compare two estimates of 3D shape: an explicit estimate from a perceptual task, requiring participants to report the perceived size of cardboard prisms through a manual matching task; an implicit estimate from a motor task, where the same individuals reached to touch the prisms with their index and thumb. The amount of depth cues available on the surface of the stimulus in each trial was varied by rotating and oscillating the prism so to hide or show disparity and motion signals and a combination thereof, according to four visual conditions (Focus, Disparity, Motion and Combined).

Explicit manual reports of 3D size were significantly affected by the amount of depth information available in a given trial (fig. 4A). Participants estimated both small and large objects with the smallest ME in the Focus condition, the largest ME in the Combined conditions, while the Disparity and Motion conditions fell in the middle, as visible in the bar graph in the inset of figure 4A. A repeated measures ANOVA on the Manual Estimations (ME) found a main effect of cue combination 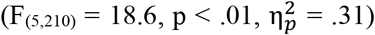, a main effect of the object size 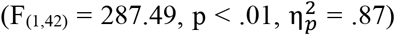 and a significant interaction between cue combination and object size 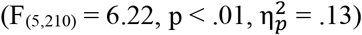.

**Figure 4.**
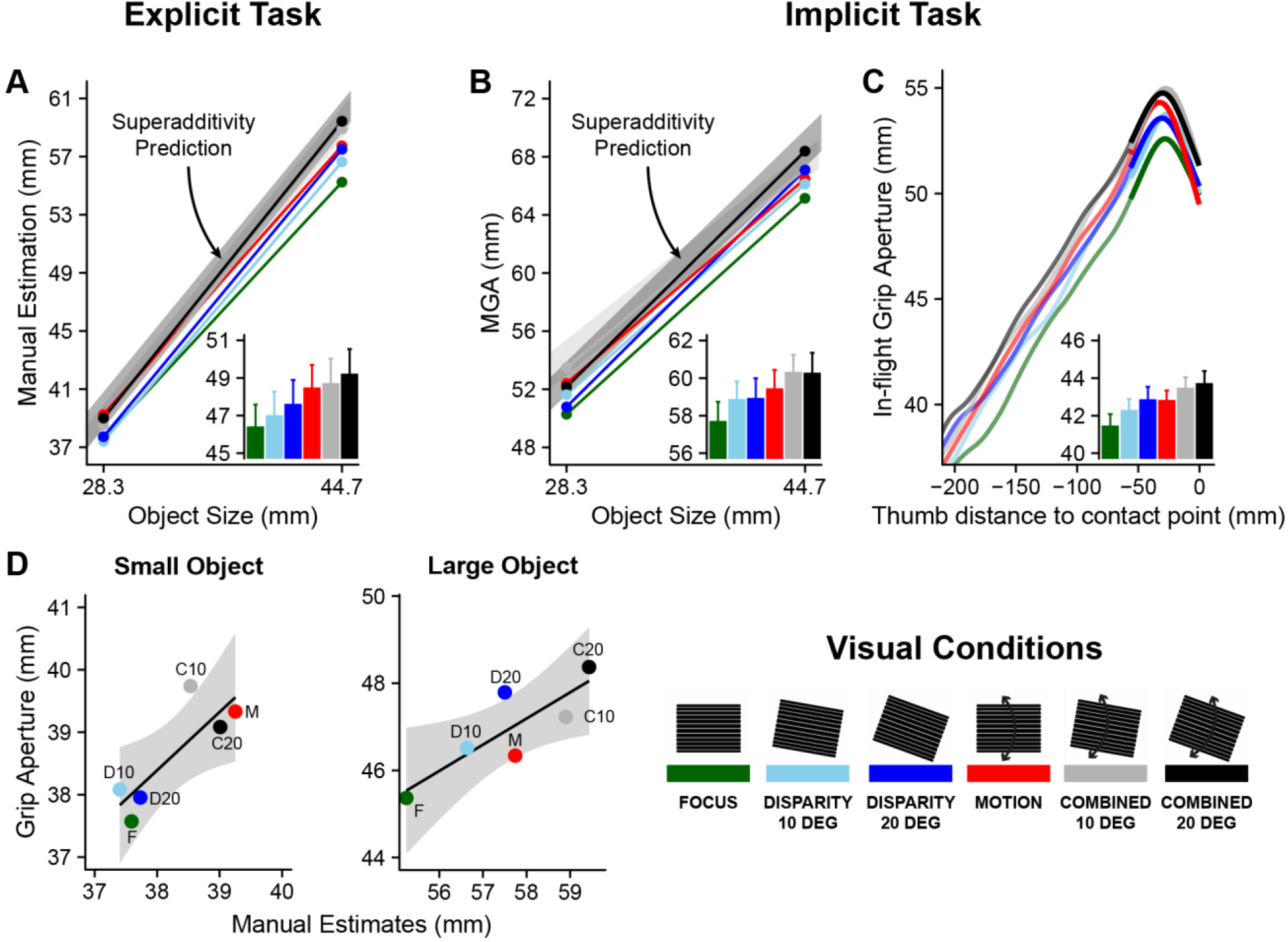
(A) Explicit 3D shape estimation: Manual Estimates as a function of the stimulus size, for each visual condition (color legend in the “Visual Conditions” panel). The inset shows the same data averaged across the stimulus size (error bars indicate between-subjects *SEM*). The gray areas represent the 95%CI of the IC model predictions for superadditivity. (B) Implicit 3D shape estimation: Maximum Grip Aperture as a function of the stimulus size, for each visual condition (data averaged across stimulus size in the inset). (C) Average Grip Aperture over the course of the reaching trajectory, as function of the instantaneous Euclidean distance of the thumb to its final contact location, for each visual condition (portions in full opacity denote effect of cue combination with p-value < .05). The inset shows the GA averaged across the entire trajectory. (D) Correlation between mean explicit reports (Manual Estimates, in abscissa) and implicit measures (mean Grip Aperture, in ordinate), for the small and the large objects (left and right panel, respectively).

A Bonferroni-Holm corrected post-hoc analysis confirmed the pattern suggested by the visual inspection of the data. The Focus condition yielded the smallest ME (smaller than the Disparity conditions averaged together: F_(1,42)_ = 12.48, p < .01; than Motion: F_(1,42)_ = 29.74, p < .01; than the Combined conditions averaged together: F_(1,42)_ = 35.71, p < .01). The average MGA in the Combined conditions was larger than in the Disparity conditions (F_(1,42)_ = 27.20, p < .01) but not larger than in the Motion condition (F_(1,42)_ = 2.63, p = .11). The ME of the Motion condition was also significantly greater than that of the Disparity condition (F_(1,42)_ = 15.65, p < .01).

The interaction term explains why the increase in depth estimation in the Combined condition relative to Motion did not reach statistical significance. An additional post-hoc analysis showed a significant difference for the large object (F_(1,42)_= 11.35, p < .01) but not for the small object (F_(1,42)_ = 1.48, p = .46). This is consistent with the IC model, which posits that the depth estimate of a given combination of depth cues is proportional to the amount of physical depth that that composite signal is estimating. As it can be seen in figure 4A, the range of the average manual estimates for the small object was less than half of the range for the large object. The small object was too small to allow the individual visual conditions to clearly spread out and separate in virtue of their different strengths, as instead happened for the large object.

The effect of tilting the object from 10 to 20 degrees was significant 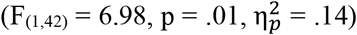 and consistent in both Disparity and Combined conditions. Overall, the ME gradually increased as more depth information was available (inset of figure 4A).

The results from the implicit task agreed almost exactly with those from the explicit task (figure 4B and 4C). This congruence in the behavior in the two cases highlights the importance that depth information has for the perception-action coupling, especially considering that even tenuous variations in the 3D structure of the scene had yet comparable effect sizes on both tasks. Participants approached both small and large objects with the smallest Maximum Grip Aperture (MGA) in the Baseline condition, the largest MGA in the Combined conditions, while the Disparity and Motion conditions fell in the middle (Fig. 4A). A repeated measures ANOVA on the MGA found a main effect of object size 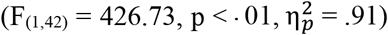, of cue combination 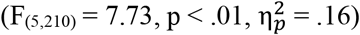, and a significant interaction between the two 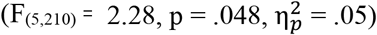.

The post-hoc results were also essentially the same. The Focus condition yielded the smallest MGA (smaller than the Disparity conditions averaged together: F_(1,42)_ = 6.65, p = .04; than Motion: F_(1,42)_ = 13.04, p < .01; than the Combined conditions averaged together: F_(1,42)_ = 19.31, p < .01). The average MGA in the Combined conditions was larger than in the Disparity conditions (F_(1,42)_ = 20.26, p < .01) but not larger than in the Motion condition (F_(1,42)_ = 2.64, p = .22). The MGA of the Disparity and Motion conditions were not different from each other (F_(1,42)_ = .84, p = .36).

A repeated measures ANOVA on a subset including the MGA of the Disparity and Combined conditions also found a significant two-way interaction between the tilt angle and the object’s size 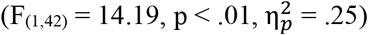, as the slope relating the MGA to the object’s size was greater when the object was tilted 20 degrees than 10 degrees. This means that the subject’s sensitivity to changes in the object’s size increased when the binocular disparities were more reliable. Again, a clearer picture of the results can be obtained by averaging across the two objects sizes, as per the inset of figure 4B, where the gradual increase of the MGA as the 3D information accumulates is most noticeable.

The Grip Aperture during the implicit task was then analyzed over the course of the entire movement, from its onset to the instant when the thumb touched the stimulus (the thumb is typically the first digit to contact an object in a front-to-back precision grasp; Volcic and Domini, 2014). A first visual inspection of the temporal evolution of the GA revealed essentially the same pattern of results found with both MGA and ME (figure 4C). This distribution was qualitatively constant throughout most of the reaching path, with a visible cue combination effect which eventually turned significant in the final part of the grasp (when the thumb was closer than 50 mm from the object).

A repeated measures ANOVA on the grip aperture averaged across the entire trajectory (inset of figure 4C) using cue combination as factor confirmed the main effect 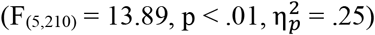. A post-hoc analysis also confirmed the pattern suggested by the visual inspection. The Focus condition yielded the smallest GA (smaller than Disparity: F_(1,42)_ = 16.75, p < .01; than Motion: F_(1,42)_ = 17.82, p < .01; than Combined: F_(1,42)_ = 49.87, p < .01). The Combined condition yielded the largest GA (larger than Disparity: F_(1,42)_ = 17.02, p < .01; than Motion: F_(1,42)_ = 13.01, p < .01). The grip apertures of the Disparity and Motion conditions were in the middle, not different from each other (F_(1,42)_ = .76, p = .39). Finally, there was a close-to-significant effect of tilting the object from 10 to 20 degrees in both Disparity and Combined conditions 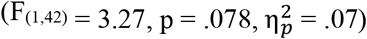.

As per the main aim of the study, the results from the Combined conditions in both explicit and implicit tasks were tested for superadditivity according to the IC model. For each condition of each task of each participant, the slope of the psychometric function relating the estimated size to the physical size was calculated through a linear fit. Each one of these slopes expressed the strength of the depth information in each condition (i.e., the term k_i_ in equation [2]). The k terms obtained from the Focus, the Disparity (10 and 20 degrees separately) and the Motion conditions were then combined according to equation [3], resulting in a prediction of the strength of the depth information in the Combined condition (k_C_). This calculation was performed for the small and large objects separately, for a total of four k_C_terms for each task (small-10deg, small-20deg, large-10deg, large-20deg). These proportionality factors were finally used to scale the physical values of the stimuli to retrieve an estimate of the stimulus size, and of the resulting psychophysical function by interpolation. Figures 4A and 4B show the 95% confidence intervals of the predicted object’s side: the light gray areas depict the predicted range for the Combined-10 condition, the dark gray areas that of the Combined-20 condition. The empirical values of both the ME and the MGA in the Combined condition were in accordance with the IC model, providing evidence in support of the idea that depth information is integrated so to maximize the sensitivity to 3D shape information. Two additional pieces of evidence further reinforce the idea that this combination rule appears to be a fundamental mechanism underlying the analysis of depth information in the brain. First, the same prediction method was successfully applied independent of the level of conscious processing of depth information, as the model predicted both the perceptual and the motor results. Second, the symmetry of the results from this study with the results of the previous experiments in VR (Campagnoli et al., 2019), demonstrate that our combination rule well describes how depth signals are translated into estimates in the visual system, and is independent of the medium (virtual versus real objects) conveying depth information to the brain.

The congruence between the perceptual and the motor performances was further assayed statistically. A 6×2 factorial ANOVA was performed crossing cue combination (6 visual conditions) with type of task (explicit or implicit). The interaction term was not significant 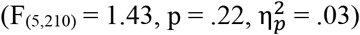, suggesting that the effect of cue combination was the same in both tasks. The ANOVA results were further confirmed by a correlation analysis between the perceptual and the motor data. For each task, the responses were averaged across each of the object sizes and each of the six viewing conditions. The datasets from the explicit task and the implicit task (average GA over the reaching) were then compared against each other. Figure 4D shows the resulting highly significant correlation that was found for both object sizes (r_smallObj_ = .84, p = .04; r_largeObj_ = .84, p = .03; n = 6).

## Discussion

When analyzing the 3D structure of an object, the visual scene provides the observer with a rich set of depth cues, ranging from monocular gradients (i.e., texture and motion) to binocular disparity. In a recent study we found that estimates of depth do not scale accurately with an object’s 3D shape, but rather they increase with the amount of depth information that specifies that shape (Campagnoli and Domini, 2019). In the present study we set out to investigate whether this behavior is the signature of a genuine superadditivity effect characterizing the processing of depth information by the visuomotor system, or instead an artifact due to top-down control caused by immersion in a virtual environment. In previous studies we proposed that the above results conform to a model of combination of depth signals termed Intrinsic Constraints model of cue integration (IC model – Domini & Caudek, 2011, 2013; Di Luca et al., 2010; Kemp et al., 2018). While the model is able to capture the behavior quantitatively, here we address the question about the nature of the phenomenon, by testing the model’s combination rule in a real-world context rather than in VR. Furthermore, to investigate a possible influence of cognition on the responses, we asked participants to report the size of cardboard prisms both through an explicit task (manual estimation) and an implicit task (reaching to touch). Participants performed both tasks under four different combinations of focus, disparity and motion depth cues. The IC model was used to predict the responses in the condition where all cues were present at the same time (Combined), based on the responses in the three intermediate conditions (Focus, Disparity and Motion).

The first significant evidence is represented by the main effect of cue combination: given the same physical prism, subjects’ estimates increased as more depth information was accessible in the scene. This pattern appeared to be independent of cognitive control, as it was detected in both explicit and implicit tasks. In the explicit task, participants increased the size of their Manual Estimates (ME) progressively from the Focus condition through Disparity, Motion and Combined. The results of the implicit task shows that this progression was inherent, systematic and robust, as it was found throughout the entire reaching trajectory and with two different object sizes. As soon as the hand was close enough to the target object, the vision of the hand allowed participants to guide their fingers onto the physical surface of the object. This is in line with previous evidence showing that what ensures a successful grasp movement is a combination of online visual control during a given trial and sensorimotor error-prediction to adjust the movements of the following trials (Bozzacchi et al., 2014, 2016; Bozzacchi & Domini, 2015; Campagnoli & Domini, 2019; Campagnoli et al., 2012; Campagnoli et al., 2015; Campagnoli et al., 2017; Cesanek et al., 2018; Cesanek & Domini, 2017; Foster et al., 2011; Volcic et al., 2013; Volcic & Domini, 2014, 2016).

The small effect size of cue combination remarks the strong interconnection between the perceptual system and the motor system during the processing of depth information. The magnitude of the effects between visual conditions was in the order of few millimeters, and yet even such subtle responses to changes in the 3D layout of the environment reverberated with similar intensity on both the grip aperture and the manual estimates.

The second major evidence is the adherence of the data to the predictions of the IC model. The model considers depth signals as scalar components of a multidimensional vector. Since in this representation each signal is assumed to be proportional to the 3D properties it encodes (e.g. depth), so it is the magnitude of the global vector. The fundamental assumption of this model is that once the scalar components have been appropriately scaled, the magnitude of the vector is the sole determinant of the perceived 3D property. It follows from this description that adding depth cues to a stimulus is equivalent to “switching on” components of this multidimensional vector with the effect of increasing its intensity, hence perceived depth. This principle can be formalized into equations predicting how depth should be estimated when multiple 3D cues are available in the scene, knowing the effect of the cues on depth estimation when acting separately. Figure 4 shows that the IC model’s predictions correctly captured the results of the Combined conditions, using the results from the Disparity, Motion and Focus conditions.

IC is therefore sufficient to explain the phenomenon of superadditivity – depth estimates increase with the amount of depth information – found both in this study and in VR (Campagnoli & Domini, 2019). Moreover, it can do so without the necessity of further ad-hoc assumptions such as prior-to-flatness (Di Luca et al., 2010; Domini and Caudek, 2003, 2009, 2010, 2011, 2013; Domini et al., 2006; Domini et al., 2011; Kemp et al., 2018).

The role of flatness information has been widely discussed in the context of a Bayesian explanation for superadditivity of disparity and texture cues. Under this framework, the combination of these particular cues has been assumed and modeled as linear, such that, if *Z*_*d*_ and *Z*_*t*_ indicate the depth estimates from disparity and texture, the combined estimate 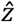 is a weighted sum 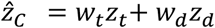 . The weights are inversely proportional to the variance of the noise associated with the depth from texture 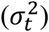 and from disparity 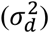 estimates and they must sum to 1. It follows that, as long as the two cues specify the same depth value *Z* = *Z*_*t*_ = *Z*_*d*_, the combined estimate is on average the same as that of each individual cue. Formally, 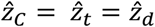 (since *w*_*t*_ + *w*_*d*_ = 1). To reconcile the empirical finding that 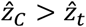 and 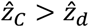, it has been hypothesized that, in virtual displays, accommodation cues and the gradient of the retinal blur are in conflict with the rest of the depth signals, in that they indicate zero depth, that of the CRT monitor (Buckley & Frisby, 1993; Frisby et al., 1995; Hoffman et al., 2008; Watt et al., 2005; Willemsen et al., 2008). Once these cues are taken into consideration, their weight can potentially affect the output of the above weighted sum. That is, if the reliability of the focus cues is sufficiently great, they are able to pull the depth estimate towards zero with greater strength when the individual depth cues are seen in isolation, relative to when the same depth cues are combined. It is therefore not surprising then that adding depth cues increases the depth estimates, because the total information about depth increases its prevalence over the flatness information carried by the focus cues. Within this framework, superadditivity should be considered as restoring flattened percepts.

Although this explanation accounts for several viewing conditions, there is one representative example case where flatness cues did not have an effect on perceived depth. In a study by Vishwanath and Hibbard (2013), observers were asked to judge the depth of a hemi-cylinder depicted through texture information. No other depth information was available on the stimulus, which was rendered on a CRT screen located at 50 cm from the observers. As expected, when the cylinder was viewed with one eye (monocular condition), the perceived depth reliably increased with the simulated depth. Most notably, depth judgements were not significantly different when the display was viewed with both eyes opened (binocular condition), which is remarkable since binocular vision introduces disparities specifying the flat surface of the screen, unlike monocular vision. Note also that a disparity field indicating a flat frontoparallel surface is highly reliable. In fact, the flatness impression derived from binocular disparity should be at least as reliable as the impression of depth conveyed by texture information, and certainly much more reliable than the flatness impression specified by the focus cues. Why did the observers not perceive the depth of the binocularly viewed cylinder as considerably flatter than the depth of the monocularly viewed cylinder? The logic of the Bayesian approach fails to explain both this phenomenon and superadditivity. This is because a critical requirement of a Bayesian estimator is that it must assess the reliability of all the sources of 3D information at all times. If focus cues are present, the estimator needs to evaluate their reliability in relation to those of the disparity and texture cues. If it detects a zero disparity field, it need to first determine whether it arises from a close-by frontoparallel surface or from an undetermined structure seen from far away. Paradoxically, this fundamental feature of the Bayesian estimator seems to work in the first case, where a faint signal flattens reliable texture and disparity estimates, explaining why combined stimuli were perceived deeper than motion or disparity stimuli. However, it falters in the second case where the signal to flatness is unmistakably reliable and should therefore dramatically reduce the perceived depth of the cylinder in Vishwanath and Hibbard (2013) study.

In contrast, the IC model can be successfully applied to both this latter case and that of superadditivity. If only disparity and texture cues are available, then, according to equation [3],

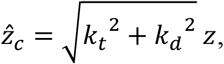

where *k*_*t*_, *k*_*d*_ are the gains associated with texture and disparity and *Z* is the simulated depth. When only disparity information is available, the gain associated with texture is nil (*k*_*t*_ = 0) and only the disparity gain contributes to perceived depth: 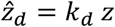. This easily explains superadditivity since 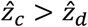. In the case of picture perception, in order to analyze this phenomenon, we need to generalize equation [3] to the case where texture and disparity stem from different depth values:

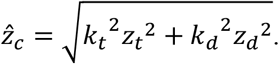

When a picture is viewed monocularly, the disparity signal is absent and therefore *k*_*d*_ = 0. When the image is viewed with two eyes, depth from disparity is zero (*Z*_*d*_ = 0). In both cases the second term under the square root vanishes and the depth estimate is solely determined by the texture signal: 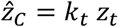 . Going back to superadditivity, note that we are not claiming that focus cues were negligible in those displays, but that they were uninfluential in both the single-cue and two-cue cases. This follows from the fact that focus cues specify the flat surface of the screen and therefore zero depth, thus adding a zero term under the square root. In summary, both the phenomenon of superadditivity and the unaltered depth perception deriving from viewing a picture with two eyes (Vishwanath, 2014) are successfully accounted for by the IC model.

The fact that our previous results obtained in VR were successfully replicated using physical objects supports the hypothesis that superadditivity is an integral feature of depth cue combination, capable to affect depth estimation across multiple layers of cognition. Furthermore, this problem taps into a more general discussion over how VR technology can be utilized as a reliable tool for research on motor control, which has received increasing attention (Freud et al., 2018; Ozana et al., 2018). The results from this study support the idea that virtual environments have the potential to be as effective as more traditional scenes, where participants see and touch real objects.

## Appendix

*Proof 1: The IC model equation maximizes the Signal-to-Noise-Ratio*

Let’s consider only two signals, for simplicity, *s*_1_ = *λ*_1_*Z* and *s*_2_ = *λ*_2_*Z. λ*_*i*_ are unknown multipliers depending on confounding variables and *Z* is the magnitude of the 3D property in question (e.g., front-to-back depth of an object). We seek an estimate 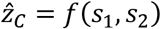 that is (1) proportional to *Z* and (2) most sensitive to 3D information while least sensitive to random fluctuations *ε*_*i*_ of *λ*_*i*_. If *λ*_*i*0_ is the unperturbed value of *λ*_*i*_: *λ*_*i*_ = *λ*_*i*0_ + *ε*_*i*_ and *s*_*i*0_ = *λ*_*i*0_*Z*. We assume small random perturbations due to changes in viewing conditions such that *ε*_*i*_ are Gaussian distributions with zero mean and standard deviations *σ*_*i*_ . Taking the derivative of 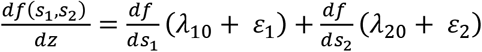, where 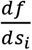 are calculated at *s*_*i0*_, we observe a signal term *S* = *f*_1_ *λ*_10_ + *f*_2_ *λ*_10_ (where 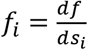) and a noise term E = *f*_1_ *ε*_1_ + *f*_2_ *ε*_2_ having standard deviation 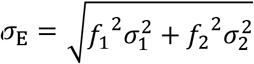 . If we minimize the Noise to Signal Ratio 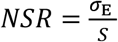 with respect to *f*_*i*_ (by solving for *f*_*i*_ the equation 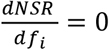) we find that the first derivatives of the sought function are 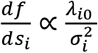 . It can be easily shown that the derivatives 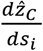 of the equation 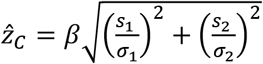 (calculated at *s*_*i0*_) meet this requirement. By substituting 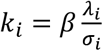 we obtain the IC equation 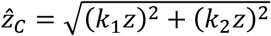 (easily generalizable to *n* signals).

*Proof 2: Derivation of* 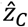 *for real stimuli*

In this study, when the stimuli were viewed under the Disparity and the Motion conditions, the total 3D shape information available resulted from the combination of each of those cues plus the focus cues. As a result, by applying equation [2], the estimated depth in the Disparity condition can be defined as:

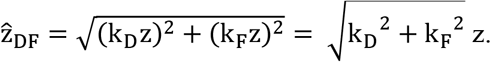

Similarly, the estimated depth in the Motion condition can be defined as:

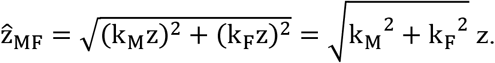

Following equation [2], in order to calculate the depth estimate in the Combined condition 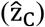 from the three sources of depth (disparity, motion and focus cues), the depth estimate from the Focus condition must be first subtracted from the depth estimates obtained in the Disparity and the Motion conditions. Equation [3] can be thus derived from equation [2] following few simple steps:

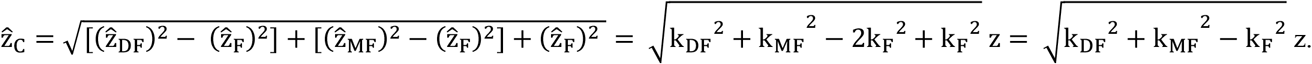

